# Enumeration of coliform bacteria and characterization of *Escherichia coli* isolated from Staff Club swimming pool in Ile-Ife, Nigeria

**DOI:** 10.1101/018127

**Authors:** Tolutope Akeju

## Abstract

Water recreation, though increasing globally, is strongly associated with infectious diseases. Unexpectedly, artificial water recreation systems e.g. swimming pools account for 90% of these outbreaks. It is therefore essential that pool waters be regularly monitored for deviations from microbial water quality guidelines. To assess the sanitary quality of a club swimming pool in Ile-Ife, Nigeria, I used the multiple-tube fermentation technique to determine the most probable number (MPN) of coliform bacteria in 100 ml of pool water. MPN estimates ranged from 9 to 93 with geometric mean of 38. *Escherichia coli* was isolated from positive presumptive tubes, indicating recent faecal contamination. The isolate elicited similar biochemical reactions as reference *E. coli* (ATCC-25922), except that it utilized sucrose and liquefied gelatin, which probably indicates potential pathogenicity. Also, the *E. coli* isolate was resistant to 13 antibiotics from 9 different classes. Finally, coliform counts and detection of *E. coli* clearly violates international guidelines. I recommend that pool operators increase water disinfection efficiency and educate the public on the need for improved swimmer hygiene to reduce the risk of recreational water illness transmission.

Acknowledgements:
The author would like to thank the Microbiology department at Obafemi Awolowo University for granting access to the Staff Club swimming pool.

**Conflicts of Interest:** The author declares no conflict of interest.

## Introduction

The recreational use of water is growing worldwide mainly because of its beneficial impact to human health (Pond 2005; WHO 2006). In the United States alone, over 301 million swimming visits were made by persons aged 7 and above in 2009 (US Census Bureau 2012). However, body-contact water recreation has been strongly associated with infectious diseases and artificial water systems – e.g. swimming pools and spas – account for more than 90% of these disease outbreaks (Doménech-Sánchez et al. 2008). Consequently, pool waters need to be monitored regularly for pathogenic microorganisms originating from faecal contamination or bather shedding e.g. *Escherichia coli* O157, *Campylobacter jejuni*, *Shigella* spp, *Cryptosporidium parvum* and Rotaviruses. Non-faecally derived pathogens like *Legionella pneumophila* and *Pseudomonas aeruginosa* have also been documented to cause recreational water illnesses (Pond 2005).

Over the years, the detection and isolation of pathogens from water have proved difficult and indicator organisms are used as surrogates. Coliform bacteria were initially used for formulating water quality standards due to their ease of enumeration via the Multiple-Tube Fermentation (MTF) technique until recent discovery about total coliforms originating from dissimilar sources (WHO 1997). While coliform genera like *Escherichia* and *Klebsiella* are mostly native inhabitants of the intestinal tract, others like *Enterobacter* and *Citrobacter* can originate from faecal, plant and soil materials (Ashbolt et al. 2001; Stevens et al. 2003). Alternatively, *E. coli* and *Enterococcus* spp provides a more reliable indication of faecal pollution and have been included as key parameters in water quality guidelines in the European Union, Australia and the US (Stevens et al. 2003).

Compared to beaches and rivers that rely on natural purification processes, the risk of disease transmission should be reduced in disinfected pool waters. Nonetheless, pool waters are highly vulnerable to swimmer-induced contamination and continuous disinfection may be unable to completely eliminate released pathogens before water ingestion (DeHaan & Johanningsmeier 1997). Research has shown that on average, adults swallow 16 ml of pool water per swimming event and children 37 ml, almost twice as adults (Dufour et al. 2006). The Centers for Disease Control and Prevention substantiated this high-exposure scenario when recent research revealed the presence of *E. coli* in 58% of pool filter samples (CDC 2013). To reduce the incidence of recreational water illnesses, microbiological quality guidelines similar to those for drinking water should apply to swimming pools.

Since brief exposures to water-borne pathogens can lead to diseases, the short-term monitoring of recreational water for deviations from microbial water quality standards is crucial to public health maintenance (WHO 2006). But water quality regulations for swimming pools or spas are yet to exist in Nigeria and scientific investigations, though scant, have consistently indicated the non-compliance of pool waters to international standards. Hence, more studies are needed to generate additional information necessary for the development of swimming pool water quality standards. This study aimed at assessing the sanitary quality of a swimming pool using total coliforms as indicators and also by characterizing any isolated organism.

## Materials and Methods

### Sampling Procedures

During a 35-day period from April to June 2009, five water samples (one per week) were collected from Staff Club Swimming Pool located at the staff quarters of Obafemi Awolowo University, Ile-Ife, Nigeria. This pool is semi-public, with bathing access restricted to only registered staff members and their guests. Pool water sampling was conducted according to standard practices (DeHaan & Johanningsmeier 1997; WHO 2006). To ensure sampling was representative of pool water quality, water was collected at a depth of 30 cm, close to swimmers and well distanced from outlets. Sampling occurred during periods of high bathing load and varied with regards to daily and weekly collection time.

### Standard Bacteriological Analysis of Pool Water

For a pool that is routinely disinfected, low coliform counts were expected and isolated organisms should provide a qualitative assessment of recent faecal contamination. To achieve this objective, the conventional MTF technique was used to determine the most probable number (MPN) of coliform bacteria present in 100 ml of pool water. This technique normally involves three steps as shown below (also see Figure 1):

**Figure 1.**
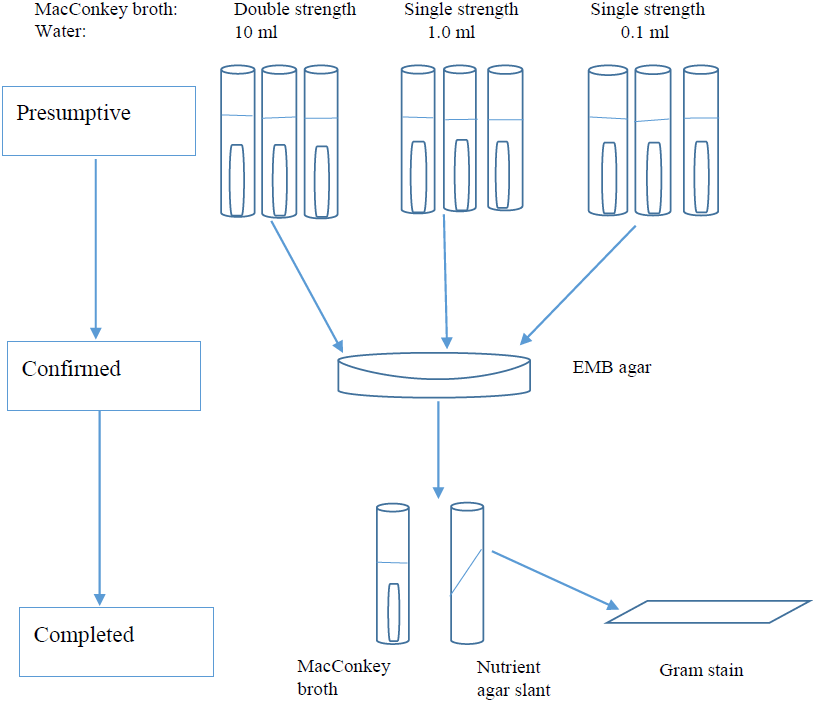
A simple illustration of the Multiple-Tube Fermentation (MTF) Technique.

#### Presumptive Test

Differential medium for the isolation of coliforms was MacConkey broth Purple. Three broth tube series – the first series containing 3 double strength broth tubes and the remaining two series comprising 6 single strength broth tubes – were inoculated with 10ml, 1ml and 0.1ml of water (ratio 3:3:3) respectively. Tubes were incubated at 37°C and observed at 24 and 48 hours. Presumptive test is positive for coliforms if acid and gas are produced in durham tubes.

#### Confirmed Test

To eliminate false-positives from non-coliform organisms, eosin methylene blue (EMB) agar plates were inoculated with a loopful from each positive presumptive broth tube by streaking across the agar surface. Plates were incubated for 24 h at 37°C.

#### Completed Test

Finally, nutrient agar slants and MacConkey broth tubes were inoculated with distinct colonies picked from cultured isolates on EMB agar plates. After incubation for 24 h at 37°C, broth cultures were observed for acid and gas production and cultured isolates on agar slants were gram stained using technique described by Aneja (2003).

### Biochemical Characterization

Besides IMViC which stands for Indole, Methyl red, Voges–Proskauer and Citrate tests, four other biochemical tests, i.e. catalase, gelatin liquefaction, starch hydrolysis and sugar fermentation were performed to confirm the identity of test isolate according to standard methods (Aneja 2003; Cheesbrough 2006).

### Antibiotic Susceptibility Test

The Kirby-Bauer disk diffusion technique was used to measure the susceptibility of the isolate to 13 commonly used antibiotics in Nigeria. Test was performed using Mueller-Hinton agar (Oxoid Code = CM0337) and two types of antibiotic multidisc (Gram positive; MICRORING/DT-NEG and Gram negative; MICRORING/DT-POS). Names, codes and concentrations of tested antibiotics are shown in Table 3.

#### Procedure

Smeared inoculum from 18-24 h nutrient broth culture of isolated organism was spread evenly on the agar surface with a sterile swab stick. Using sterile forceps, antibiotic multidiscs were placed at the center of inoculated media. Plates were inverted and incubated at 37°C for 24 h. Thereafter, zones of inhibition around the discs were observed, their diameters measured and classified as resistant (R), susceptible (S) or intermediate (I) according to interpretive criteria defined by the Clinical and Laboratory Standards Institute (CLSI 2007).

### Statistical Analysis

MPN of coliform bacteria per 100 ml of original water sample was determined from a standard 3-tube statistical table (WHO 1997). Geometric mean (Geomean), standard deviation (SD) and coefficient of variation (CV %) were calculated using predefined functions in MS Excel. Equation for the geometric mean is stated below:

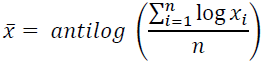

## Results

Change in broth colour from purple to yellow and presence of gas in durham tubes indicated the presence of coliforms in positive presumptive tubes. On EMB agar, confirmed test for coliform bacteria showed the appearance of *E. coli* alone – distinguished by the typical greenish metallic sheen. In the completed tests, acid and gas were observed in broth tubes and gram stain revealed red, non-spore forming rods, indicating the bacterial isolate is gram-negative. The MPN of coliform bacteria, as presented in Table 1, ranged from 9 to 93. Geomean, SD, and CV% of MPN values were 38, 2.37 and 6.3% respectively. The 95% confidence limits (Geomean ± 2SD) were 6.7 (lower limit) and 211.6 (upper limit).

**Table 1.**
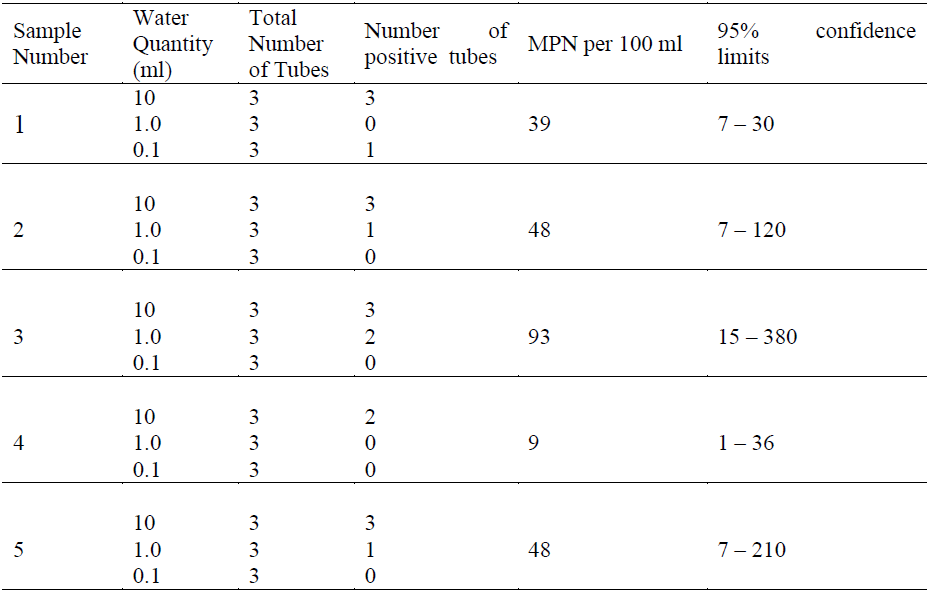
Most Probable Number of coliform bacteria in staff club swimming pool, OAU, Ile-Ife.

In Table 2, comparisons of biochemical reactions for the reference *E. coli* strain (ATCC 25922) as outlined by Siegrist (2011) and the *E. coli* isolate showed differences in gelatin liquefaction and sucrose fermentation reactions. Antibiotic susceptibility tests for the *E. coli* isolate revealed resistance to all tested antibiotics (see Table 3).

**Table 2.** Biochemical characteristics of *E. coli* isolated from Staff Club Swimming Pool.

| Test | Reaction |  |
| --- | --- | --- |
|  | Isolated *E. coli* | Reference *E. coli*<br>(ATCC 25922) |
| Indole | + | + |
| Methyl red | + | + |
| Voges-Proskauer | — | — |
| Citrate | — | — |
| Catalase | + | + |
| Starch hydrolysis | — | — |
| Gelatin liquefaction | + | — |
| Mannitol | + | + |
| Glucose | + | + |
| Sucrose | + | — |
| Lactose | + | + |
| Inositol | — | — |

**Table 3.** Antibiotic susceptibility profile of *E. coli* isolated from Staff Club Swimming Pool.

| Disc | Antibiotic |  |  | Inhibition diameter (mm) | Interpretation |
| --- | --- | --- | --- | --- | --- |
|  | Name | Code | Concentration (µg) |  |  |
| Gram-Positive | Streptomycin | STR | 25 | 9.0 | R |
|  | Tetracycline | TET | 25 | 0 | R |
|  | Colistin | COL | 25 | 0 | R |
|  | Gentamycin | GEN | 10 | 7.5 | R |
|  | Nalixidic acid | NAL | 30 | 0 | R |
|  | Ampicillin | AMP | 25 | 0 | R |
|  | Nitrofurantoin | NIT | 200 | 0 | R |
|  | Cotrimazole | COT | 25 | 0 | R |
| Gram-Negative | Streptomycin | STR | 10 | 8.5 | R |
|  | Erythromycin | ERY | 5 | 0 | R |
|  | Penicillin | PEN | 11 | 0 | R |
|  | Tetracycline | TET | 10 | 0 | R |
|  | Gentamycin | GEN | 10 | 8.0 | R |
|  | Cloxacillin | CXC | 10 | 0 | R |
|  | Chloramphenicol | CHL | 10 | 0 | R |
|  | Ampicillin | AMP | 10 | 0 | R |

## Discussion

### Validity of laboratory test procedures

Low co-efficient of variation (6.3%) coupled with the fact that range of MPN estimates (9 – 93) was contained within 95% confidence limits (6.7 – 211.6) of the geometric mean validates intra-laboratory test procedures. However, the 95% confidence intervals of the individual coliform counts (see Table 1) and the geometric mean are wide enough to reveal the inherent low precision of MPN estimates. Ever since, researchers have duly recommended increasing the number of tube replicates and samples to overcome this imprecision and compared to plate counts, the MPN can provide more accurate estimates when bacteria counts are low (Sutton 2010).

### Are total coliforms suitable indicators of pool water quality?

Recently, inclusion of total coliforms in compliance testing has been strongly debated due to their heterogeneous origins (Ashbolt et al. 2001; Stevens et al. 2003). Even the WHO (2006) excluded total coliforms from among suitable microbial parameters in the guidelines for safe recreational water. Nevertheless, studies have repeatedly shown that even ‘true faecal indicators’ are unlikely to correlate with pathogen densities in water at low pollution levels (Payment & Locas 2011). Additionally, densities of faecal coliforms and faecal streptococci in pool waters are usually low, making them unsuitable indicators (Seyfried 1989). But total coliforms are present in sufficient densities, sensitive to chlorination and therefore reliable for assessing the efficiency of sanitary processes such as the disinfection of swimming pool waters (Ashbolt et al. 2001; Nikaeen et al. 2009).

### Assessment of estimated MPN of coliform bacteria in Staff Club Swimming Pool

Apart from WHO (2006) guidelines which state that ‘thermo-tolerant’ coliforms or *E. coli* should be below 1/ 100 ml, several national organizations have bacteriological standards for pool waters that are more or less similar. According to German (DIN 19643/1984), British (BSI PAS 39:2003) and Greek (443/B/1974) regulations, total coliform counts should not exceed 0, 10 and 14 per 100 ml and *E. coli* should be totally absent in 100 ml of pool water (Papadopoulou et al. 2008). Therefore, when assessed in the light of aforementioned regulations, the coliform bacteria counts estimated in this study were clearly above recommended limits and the bacteriological quality of this pool can be deemed unacceptable.

### Biochemical characterization of *E. coli* isolate

Here, I discuss the significance and implications of observed differences in the biochemical profiles of *E. coli* and reference *E. coli* ATCC. The isolated *E. coli* can be identified as biotype I based on IMViC reaction pattern (+ + − −) (Odonkor & Ampofo 2013). In addition, knowing that most wild-type strains of *E. coli* are unable to produce α-amylase for starch hydrolysis (Rosales-Colunga & Martínez-Antonio 2014) and that only ≤ 10% of commensal and pathogenic *E. coli* strains can ferment inositol (Leclercq et al. 2001) can help explain the negative reactions obtained for starch and inositol respectively.

Most industrial *E. coli* strains are unable to liquefy gelatin and gelatinase is a virulence factor moderately expressed in pathogenic *E. coli*. For example, one study in South Africa reported gelatinase production in 60% of verotoxic *E. coli* isolates from water and wastewater samples (Doughari et al. 2011). In addition, while 19.4% of *E. coli* isolates from urinary tract infection patients produced gelatinase, none from healthy persons was gelatinase positive (Shruthi et al. 2012). Therefore, the gelatin liquefaction elicited by this *E. coli* isolate may indicate pathogenicity. Sucrose-utilizing *E. coli* strains are mostly pathogenic, and *E. coli* W (ATCC 9637) is the only sucrose positive commensal/laboratory strain (Sabri et al. 2013). Because most pathogenic *E. coli* strains belong the biotype I group, the positive reaction for sucrose may infer pathogenicity but serological and molecular testing would be needed for confirmation (Leclercq et al. 2001; Odonkor & Ampofo 2013).

### Antibiotic resistance

The *E. coli* isolate was resistant to all 13 antibiotics, which is unsurprising because of the rise of antimicrobial resistance. The WHO (2014) noted that though antimicrobial resistance is a ‘normal evolutionary process’, the widespread and indiscriminate use of antibiotics in human and veterinary medicine have escalated this process in recent decades (WHO 2014). In the US for instance, resistance of *E. coli* isolates to ≥ 3 classes of antibiotics (i.e. multi-drug resistance) increased from 7% in the 1950s to 64% in the 2000s (Tadesse et al. 2012).

*E. coli* acquires resistance genes easily and has recently shown resistance, not just to older, commonly used antibiotics, but also to fluoroquinolones and third generation cephalosporins (Tadesse et al. 2012; WHO 2014). Multi-drug resistance (MDR) in *E. coli* is probably much worse in developing countries (Collignon 2009) and in Nigeria for example, resistance of *E. coli* isolates from students to tetracycline, ampicillin, chloramphenicol and streptomycin increased from (9 – 35)% in 1986 to (56 – 100)% in 1998 (Okeke et al. 2000).

Since the *E. coli* isolated in this study is most probably derived from swimmers, human origin is presumed and MDR in human isolates is usually high. Consider one study on the susceptibility of 128 *E. coli* isolates to 13 antibiotics. *E. coli* isolates derived from humans were resistant to 2–13 antibiotics with mean resistance index of 0.67, four times greater than resistance index (0.17) for animal isolates which were resistant to just 1–6 antibiotics. (Vantarakis et al. 2006). In contrast, another study revealed higher antibiotic resistance in *E. coli* isolates from ‘food animals’. Specifically, 59.1% of *E. coli* isolates from cattle, 53.7% from pigs and 55.1% from chicken exhibited MDR, compared to just 19.5% isolates from humans. Also, from the 796 pan-susceptible *E. coli* isolates, 80% were from humans, 8.7% from cattle, 7.5% from pigs and 3.9% from chickens (Tadesse et al. 2012). Perhaps, these two findings do not contrast so sharply. Indications have recently emerged that MDR *E. coli* strains can be transferred to humans through the food chain. The consumption of animal meat – particularly poultry where antibiotics are frequently used in feed – is the most likely source of multi-resistant *E. coli* in humans (Collignon 2009).

## Conclusions

The unacceptable bacteriological pool water quality in this study indicates an increased risk for the transmission of RWIs. Besides enforcing adequate disinfection levels and compliance to microbial standards, pool operators must educate swimmers on the need for improved hygiene practices to prevent RWIs. For example, pre-swim showers, regular bathroom breaks, and not swimming during a gastro-enteric illness can significantly reduce the amount of urine, sweat and faecal material introduced into pool waters (CDC 2013). It must be noted that pool water quality assessments using only total coliforms and *E. coli* may be inadequate. For robust water quality assessments, monitoring for chemical parameters and non-faecally derived bacteria e.g. *Legionella* and *P. aeruginosa* is recommended (WHO 2006). High MDR in isolated *E. coli* highlights the growing threat of antibiotic resistance. Since no new class of antibiotics have been discovered since the 1980s (WHO 2014), public health efforts must be geared towards curbing the spread of antibiotic resistance, while the search for novel antimicrobials continue.

